# Divalent ligand-monovalent molecule binding

**DOI:** 10.1101/2021.01.14.426724

**Authors:** Mathijs Janssen, Harald Stenmark, Andreas Carlson

## Abstract

Simultaneous binding of divalent ligands to two identical molecules is a widespread phenomenon in biology and chemistry. Here, we describe this binding event as a divalent ligand AA that can bind to two identical monovalent molecules B to form the complex AA · B_2_. Cases where the total concentration [AA]_*T*_ is either much larger or much smaller than the total concentration [B]_*T*_ have been studied earlier, but a description of intermediate concentrations is missing. In this paper, we describe the general case of any ratio of *ξ* ≡ [B]_*T*_ /[AA]_*T*_. We show that the concentration of the intermediate complex AA · B is governed by a cubic equation and discuss several scenarios in which this cubic equation simplifies. Our numerical results, which cover the entire range of 0 < *ξ* < ∞, are relevant to processes wherein the concentrations of free ligands and proteins both decrease upon binding. Such ligand and protein depletion is expected to be important in cellular contexts, e.g., in antigen detection and in coincidence detection of proteins or lipids.

## I. INTRODUCTION

Chemical binding is at the heart of many processes in biology, including oxygen binding to hemoglobin, self assembly, antibodies binding to antigens, and growth factors binding to their transmembrane receptors [1–6]. In many cases, binding interactions should be specific and strong, yet reversible [7–10]. One way to accomplish such a “molecular velcro” [7] is through ligands containing many ligating units per molecule: Multivalent ligands are known to bind transmembrane receptors more readily than their monovalent counterparts (with one binding site per ligand). This makes multivalent ligands interesting in clinical applications, for example, where less therapeutic cargo is needed for the same response. The intuitive explanation why multivalent ligands bind more readily to, for instance, receptors on a plasma membrane or a viral envelope, is that, after the binding of a first ligating unit with association constant *K*_1_, the other ligating units are close to other membrane-bound receptors as well. Around a first bound unit, a second ligating unit is thought to sweep out a semi circle with a radius set by the (fixed) distance between ligating units [11–14]. This is typically a nanometers length, meaning that the *effective concentration* of ligating units belonging to a partly-bound multivalent ligand is much higher than the concentration of unbound ligands nearby. More generally, for flexible rather than stiff linkers between ligating units [15, 16], increased effective concentrations can be determined rigorously by statistical mechanics [17, 18].

In turn, high effective concentrations are reflected in a high association constant *K*_2_ for binding a second ligating unit of a multivalent ligand, and the same for further binding steps. Systems for which *K*_2_/*K*_1_ > 1 are called *cooperative* [19–22]. In the above example of large effective concentrations, one speaks of apparent cooperativity. This is to distinguish it from true cooperativity, which refers to binding pockets whose binding affinity changes when nearby pockets are occupied, as happens for the binding of oxygen to hemoglobin [23]. In either way, the hallmark of cooperativity is the switching from mostly-unbound to mostly-bound ligands over a narrow protein-concentration range [19].

Ligand-protein binding models often have governing equations that simplify when one molecular species is assumed to be present in excess compared to other species. While this assumption may be appropriate to certain systems and experiments, it is not always the case. One example is when two types of ligands compete for the binding of one type of receptor. In this case, the relative concentrations of the ligands must be important—unless the receptor is in excess to both types of ligand, in which case there would be no competition for it. When no molecular species is in excess to the other present in the system, binding can significantly reduce the concentration of unbound species. Such depletion is difficult to capture in theoretical models, even for the steady state, as governing algebraic equations are typically nonlinear and with a high polynomial order. Two notable exceptions where the concentrations of all species can be expressed analytically are monovalent ligand-monovalent receptor binding [1] and the competitive binding of two different monovalent ligands to one type of monovalent protein [24].

Several recent review articles [19, 20, 22] discuss the reversible binding of a divalent ligand AA to two identical monovalent proteins B [Fig. 1(a)],

**Figure 1.**
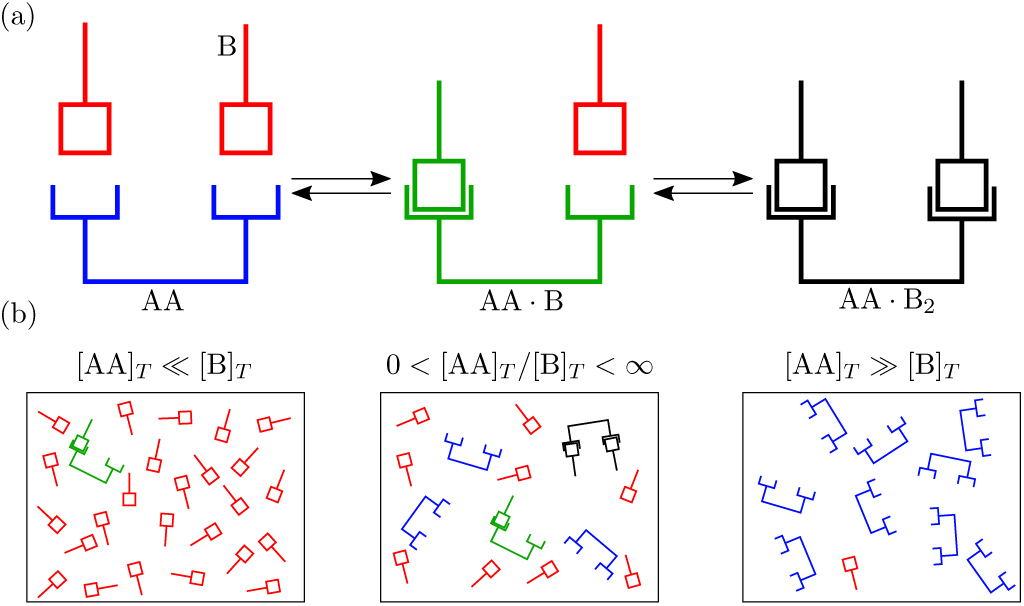
(a) Binding of two monovalent proteins B to a divalent ligand AA, to form the complexes AA · B and AA · B_2_. (b) Different relative concentrations of [AA]_*T*_ and [B]_*T*_.

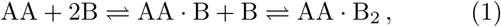

as it is the simplest example of a binding reaction with nontrivial effects of multivalency and cooperativity. Yet, Eq. (1) has value in its own right: It captures hormone action [25] and the binding of divalent antibodies to antigens on pathogens [11, 12, 26–30]. Moreover, reaction of the type of Eq. (1) were also realized in synthetic systems

From the law of mass action follow the reaction-rate equations associated with Eq. (1). In turn, the steady state of these equations yields two association constants *K*_1_ and *K*_2_ as [see Appendix A]

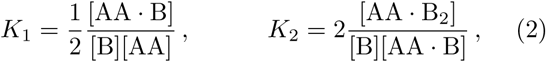

where factors of 1/2 and 2 account for the degeneracy of the intermediate complex AA · B. An assumption underlying the derivation of Eq. (2) in terms of concentrations, is that all species are well mixed. This assumption may be violated when receptors cluster at the plasma membrane [31, 32].

The reaction in Eq. (1) does not affect the total concentration [AA]_*T*_ and [B]_*T*_ of ligands AA and proteins B—both bound and unbound—and, hence,

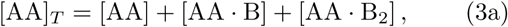

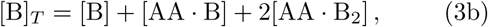

needs to be satisfied.

From the four expressions in Eqs. (2) and (3), in principle, the four unknown concentrations ([AA], [B], [AA · B] and [AA · B_2_]) can be determined. Perelson and DeLisi [29] studied the same equations for [AA]_*T*_ ≫ [B]_*T*_ by asserting [AA]_*T*_ [AA]; Hunter and Anderson [19] studied the same equations for [AA]_*T*_ ≪ [B]_*T*_ by asserting [B]_*T*_ ≈ [B] [Fig. 1(b)]. As we move away from these limits, neither [AA]_*T*_ ≈ [AA] nor [B]_*T*_ ≈ [B] will hold as the binding reaction in Eq. (1) causes ligand and protein depletion. Here, we study Eqs. (2) and (3) over the complete range 0 < [B]_*T*_ /[AA]_*T*_ < ∞.

## II. MODEL

Inserting Eqs. (3a) and (3b) into Eq. (2) yields [AA · B] = 2*K*_1_ ([B]_*T*_ − [AA · B] − 2[AA · B_2_])

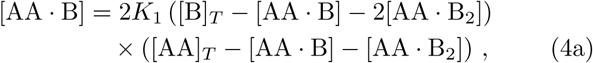

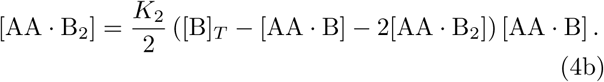

Next, we rewrite Eq. (4) in terms of the dimensionless concentrations *x*_3_ = [AA · B]/[AA]_*T*_ and *x*_4_ = [AA · B_2_]/[AA]_*T*_, with the dimensionless association constants (or, equivalently, “normalized concentration” scales [19]) *κ*_1_ = *K*_1_[B]_*T*_ and *κ*_2_ = *K*_2_[B]_*T*_, and with the ligand-to-receptor ratio *ξ* = [B]_*T*_ /[AA]_*T*_,

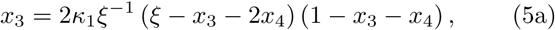

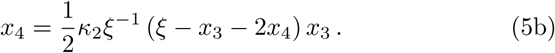

We rewrite Eq. (5b) to

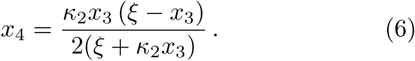

Inserting Eq. (6) into Eq. (5a) yields

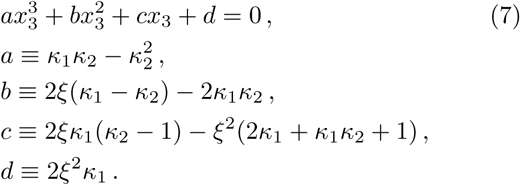

While Eq. (7) for *x*_3_ can be solved analytically with Cardano’s formula, unfortunately, its solution, presented in Appendix B, is too cumbersome to be helpful. In Appendices C–E, we analyse Eq. (7) for limiting values of the ligand-to-receptor ratio, *ξ* 1 and *ξ* 1, and for the case where the cooperativity parameter *α* = *K*_2_/*K*_1_ takes the value *α* = 1. The analytical results obtained there help us interpret the numerical solutions of Eq. (7) that we present below.

## III. RESULTS

After Eq. (4), we reduced the four parameters [AA]_*T*_, [B]_*T*_, *K*_1_, and *K*_2_ of our original problem [Eqs. (2) and (3)] to three dimensionless combinations *κ*_1_, *κ*_2_, and *ξ* thereof. We choose these particular combinations to tidy up the calculations of Section II and Appendices B–F. But for the description of particular systems or experiments, other dimensionless combinations of the four dimensional parameters may be more appropriate. Accordingly, to recover the results of Ref. [29], we first vary *K*_1_[AA]_*T*_ for several *ξ*, fixing either *K*_2_[AA]_*T*_ or *K*_2_[B]_*T*_:

Figure 2 shows [AA · B]/[AA]_*T*_ (a) and [AA · B_2_]/[B]_*T*_ (b) as obtained from Eqs. (7) and (6) as a function of *K*_1_[AA]_*T*_ for *ξ* = {0.2, 1, 1.5, 2, 5, 100} and *K*_2_[AA]_*T*_ = 1 (a) and *K*_2_[B]_*T*_ = 1 (b). Here, panel (b) generalizes the “cross linking curves” of Fig. 3 of Ref. [29] to *ξ* values away from the limit *ξ* → 0. For *ξ* = 0.2, we show Eq. (D5) (purple dashed line) as obtained by Ref. [29]. Small difference are visible in Fig. 2(b) between the purple dashed line and the purple diamonds, which means that, for *ξ* = 0.2, Eq. (D5) approximates the numerical solution to Eqs. (6) and (7) well, but not perfectly. This finding is in line with Appendix D, where we find that Eq. (D5) contains errors of 𝒪(*ξ*^3^). Reference [29] showed that max 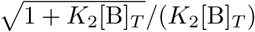, In our case of *K*_2_[B]_*T*_ = 1, we find max([AA · B_2_]/[B]_*T*_) = 0.0857, which is indeed observed in Fig. 2(b) up to *ξ* = 1. Moreover, the bell shape of Eq. (D5) was shown to be symmetric around its maximal value [29]. With increasing *ξ*, however, we see that this symmetry is broken. For *ξ* > 2, [AA · B_2_]/[AA]_*T*_ and [AA · B_2_]/[B]_*T*_ are sigmoidal instead. Next, we show Eq. (C2) (thick grey line), which corresponds to the limit *ξ* → ∞ [19]. Small differences between this expression and the numerical solution to Eqs. (6) and (7) for *ξ* = 100 are visible in both panels of Fig. 2. This finding is in line with Appendix C, where we find that Eq. (C2) contains errors of 𝒪(*ξ*^−1^). Finally, we note that bell-shaped [AA · B]-curves appear for varying *K*_1_[AA]_*T*_ only if one fixes either *K*_2_[AA]_*T*_ or *K*_2_[B]_*T*_. In experiments, concentrations are often more easily varied than association constants. Yet, varying [AA]_*T*_ at fixed *K*_1_ and *K*_2_[AA]_*T*_ would require *K*_2_ to vary as ∼ 1/[AA]_*T*_.

**Figure 2.**
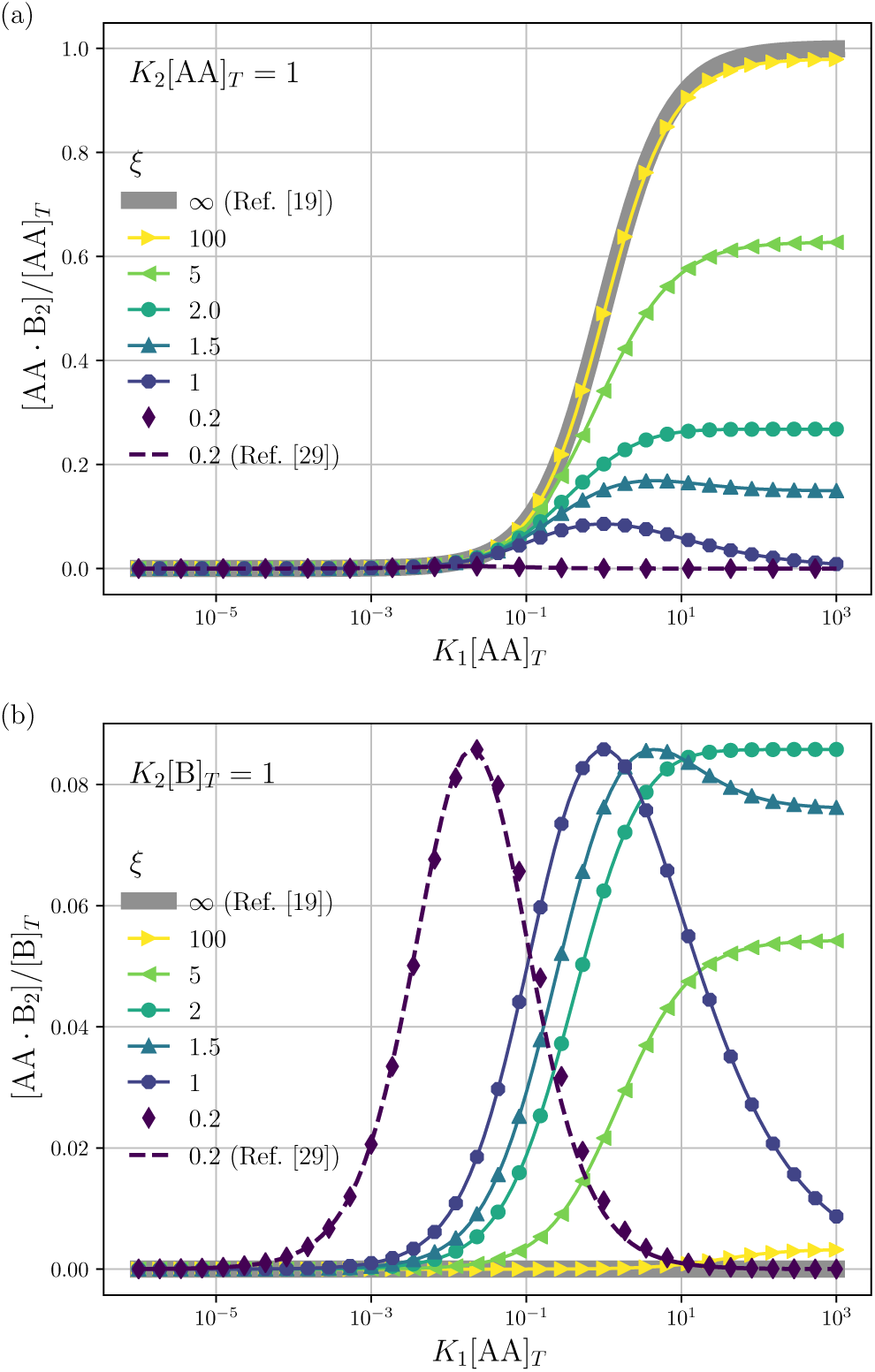
We show [AA · B_2_]/[AA]_*T*_ (a) and [AA · B_2_]/[B]_*T*_ (b) as a function of *K*_1_[B]_*T*_ for *K*_2_[AA]_*T*_ = 1 (a) and *K*_2_[B]_*T*_ = 1 (b) and several *ξ* ≡ [B]_*T*_ /[AA]_*T*_. As in Perelson and DeLisi [29], [AA · B_2_]/[B]_*T*_ has a bell shape for small *ξ*.

**Figure 3.**
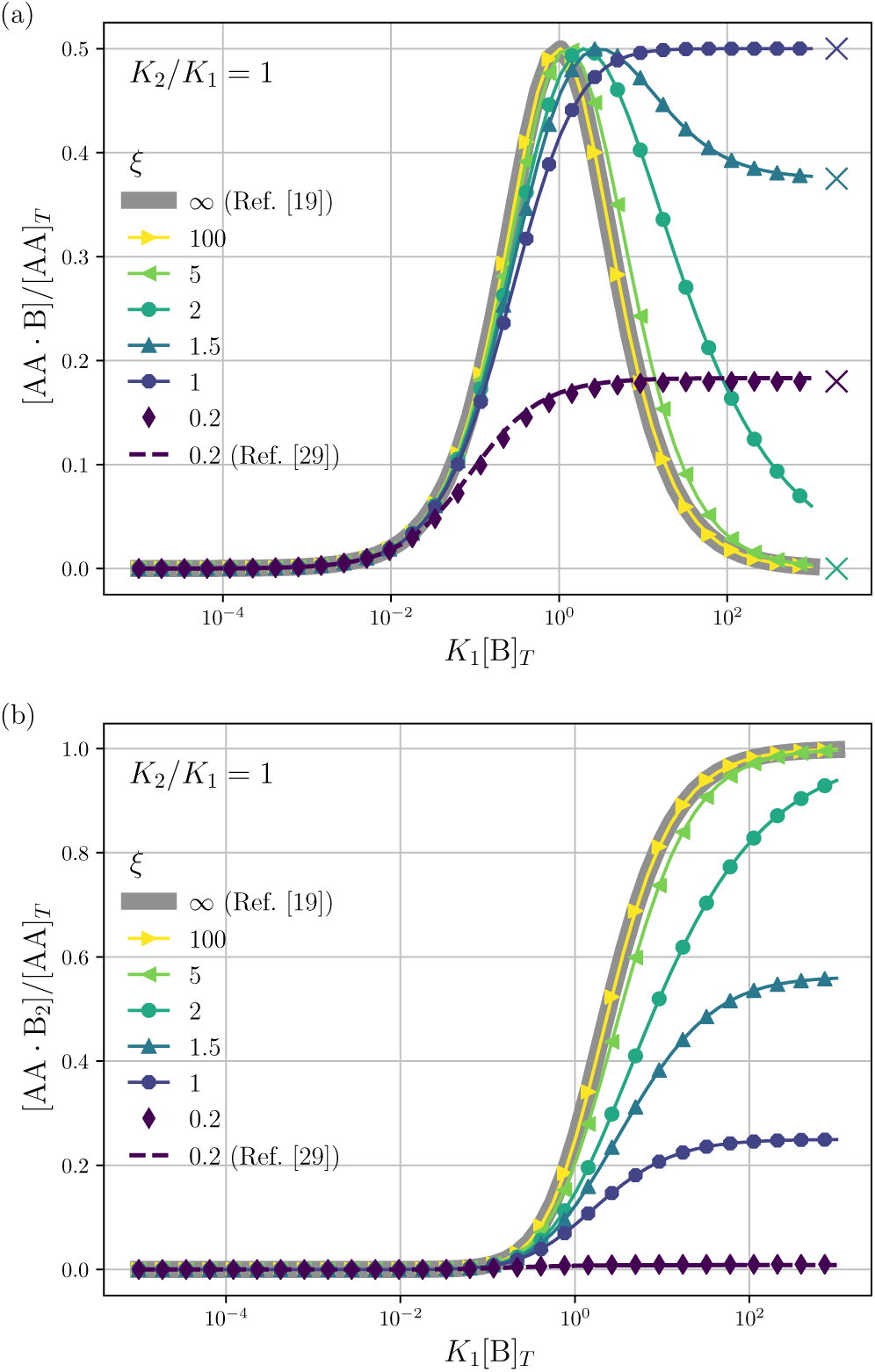
Theoretical predictions for a dilution experiment. We show [AA · B]/[AA]_*T*_ (top row), [AA · B_2_]/[AA]_*T*_ (middle row) as a function of *K*_1_[B]_*T*_ for *α K*_2_/*K*_1_ = 1 for *ξ* ≡ [B]_*T*_ /[AA]_*T*_ = 0.2, 1, 1.5, 2.0, 5.0. Also shown are approximations to [AA · B]/[AA]_*T*_ and [AA · B_2_]/[AA]_*T*_ for *ξ* ≫1 [see Eqs. (C1)–(C3) and Ref. [19]] and for *ξ* ≪ 1 [see Eq. (D4) and Ref. [29]]. Panel (a) shows the analytical predictions from Eq. (E4) for *K*_1_[B]_*T*_ ≫ 1 with crosses.

Next, to describe a dilution experiment, we vary [AA]_*T*_ and [B]_*T*_ at fixed *ξ* = [B]_*T*_ /[AA]_*T*_. Different from before, we fix the dimensionless cooperativity parameter *α* ≡ *K*_2_/*K*_1_, as it is often set solely by (fixed) molecular properties [17, 18]. Figure 3 shows numerical results for [AA · B]/[AA]_*T*_ (a) and [AA · B_2_]/[AA]_*T*_ (b) as a function of *κ*_1_ = *K*_1_[B]_*T*_, for several *ξ* and *α* = 1. In this case (*α* = 1), the cubic term in Eq. (7) vanishes, and the remaining quadratic equation can be easily solved analytically [cf. Eq. (E2)]. Moreover, [AA · B_2_]/[AA]_*T*_ is governed by a simple expression [Eq. (E5a)]. A salient feature of the curves in Fig. 3(a) are the plateaus for *K*_1_[B]_*T*_ ≫ 1. For *ξ* ∼ 1, their height is given by 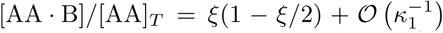 [cf. Eq. (E4)], as indicated by the crosses in Fig. 3(a). Notably, the maximal plateau height occurs at *ξ* = 1, as also follows from Eq. (E4). Next, we compare our numerical results for *ξ* = 0.2 (purple diamonds) to the expressions derived in Ref. [29] [cf. Eqs. (D4) and (D5)] (purple dashed lines). These panels reinforce our analytical insights of Appendix D, namely, that the expressions derived in Ref. [29] contain errors of 𝒪(*ξ*^3^); hence, describe [AA · B]/[AA]_*T*_ and [AA · B_2_]/[AA]_*T*_ decently, but not perfectly, at *ξ* = 0.2. Finally, note that [AA · B] cannot exceed the total concentrations of its constituents, [AA]_*T*_ and [B]_*T*_; hence, 0 < [AA · B]/[AA]_*T*_ < min(1, *ξ*). Likewise, for [AA · B_2_], we find that 0 < [AA · B_2_]/[AA]_*T*_ < min(1, *ξ*/2). The data in Fig. 3 satisfies these constraints. For the same parameters as in Fig. 3, Fig. 4 shows the receptor occupancy *θ* ≡ *x*_3_/2 + *x*_4_ = {[AA · B]/2 + [AA · B_2_]} /[AA]_*T*_ for *α* = 1 [Fig. 4(a)] and *α* = 100 [Fig. 4(b)]. We see that increasing cooperativity shifts *θ* curves to smaller *K*_1_[B]_*T*_ values and that *θ* switches from *θ* ≈ 0 to *θ* ≈ 1 over a narrower range of *K*_1_[B]_*T*_. To characterise the slope of *θ*, we numerically determined the Hill coefficient

**Figure 4.**
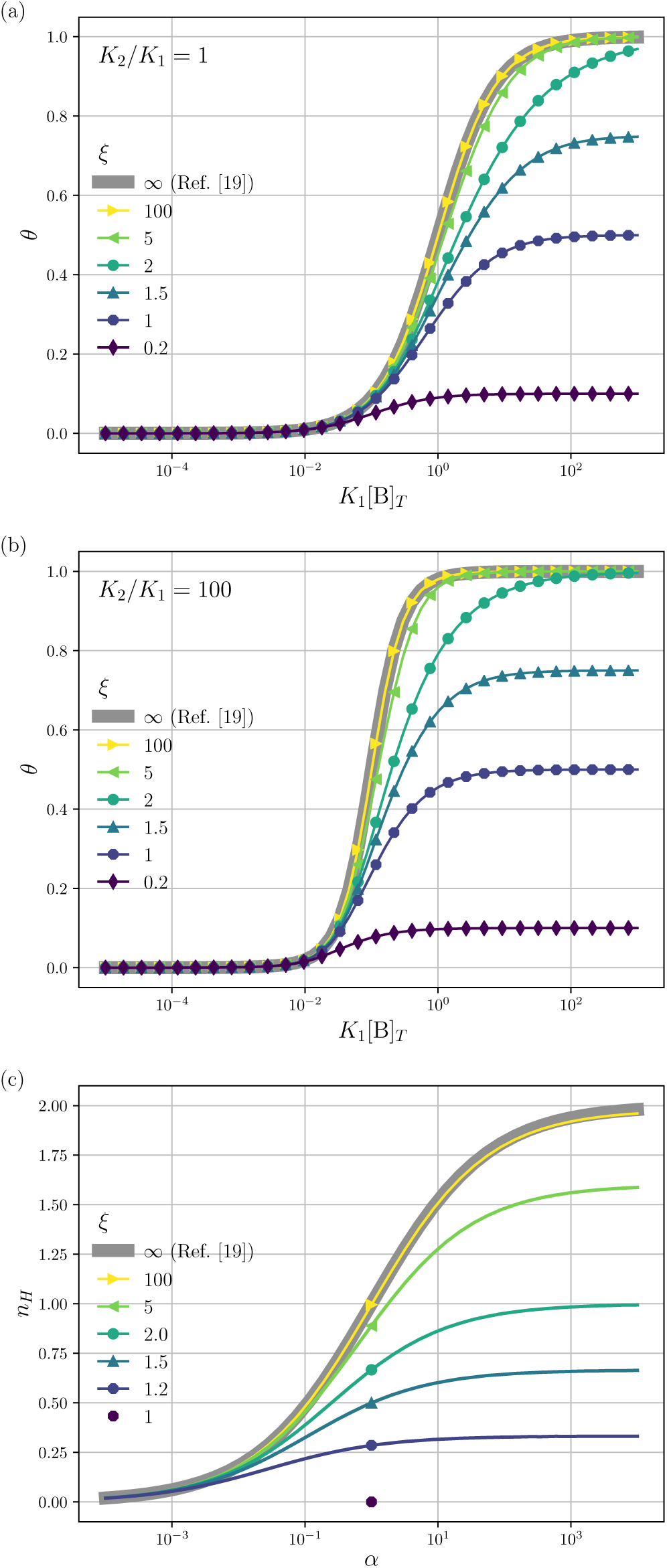
The receptor occupancy *θ* for *α* = 1 (a) and *α* = 100 (b) and other parameters as in Fig. 3. Panel (c) shows the Hill coefficient *n*_*H*_ [Eq. (8)] for several *ξ* ≥ 1 (lines). The thick grey line shows Eq. (C4), corresponding to *ξ* → ∞. We also show the predictions of Eq. (E7) for *α* = 1 (dots).

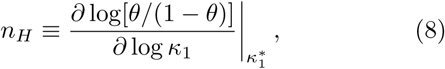

where 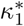 is such that 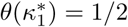; hence,

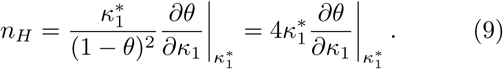

Figure 4(c) shows the *α* dependence of *n*_*H*_ for several *ξ*. As such, this figure generalizes Fig. 6 of Ref. [19], which showed *n*_*H*_ for *ξ* ≫ 1 [Eq. (C4)], indicated here with a thick grey line. We see that, for *ξ* = 100, the numerically determined *n*_*H*_ is close to predictions from Eq. (C4). Conversely, we see that *n*_*H*_ → 0 for *ξ* → 1. For *ξ* < 1, we see in Fig. 4 (a) and (b) that *θ* < 1/2, leaving *n*_*H*_ undefined. The dots in Fig. 4(c) for *α* = 1 represent the analytical expression Eq. (E7), which gives a perfect match when compared with the numerical prediction.

At last, we mimic a titration experiment by varying [B] at fixed *K*, *K* and [AA], *i*.*e*., varying *κ*_1_ at fixed *α* and *K*_1_[AA]_*T*_ (or *κ*_2_/*ξ*, in terms of our original dimensionless parameters). Figure 5 shows [AA · B]/[AA]_*T*_ (a) and [AA · B_2_]/[AA]_*T*_ (b) for *α* = 10 and various *K*_1_[AA]_*T*_. For *K*_1_[AA]_*T*_ = 1/10, we see that [AA · B]/[AA]_*T*_ and [AA · B_2_]/[AA]_*T*_ are similar to the curves of Fig. 3 for *ξ* ≫ 1. Different from Fig. 3(a) and (b) is that [AA · B]/[AA]_*T*_ does not develop plateaus at large *K*_1_[B]_*T*_. Instead, both [AA · B]/[AA]_*T*_ and [AA · B_2_]/[AA]_*T*_ shift towards larger *K*_1_[B]_*T*_ for larger *K*_1_[AA]_*T*_.

**Figure 5.**
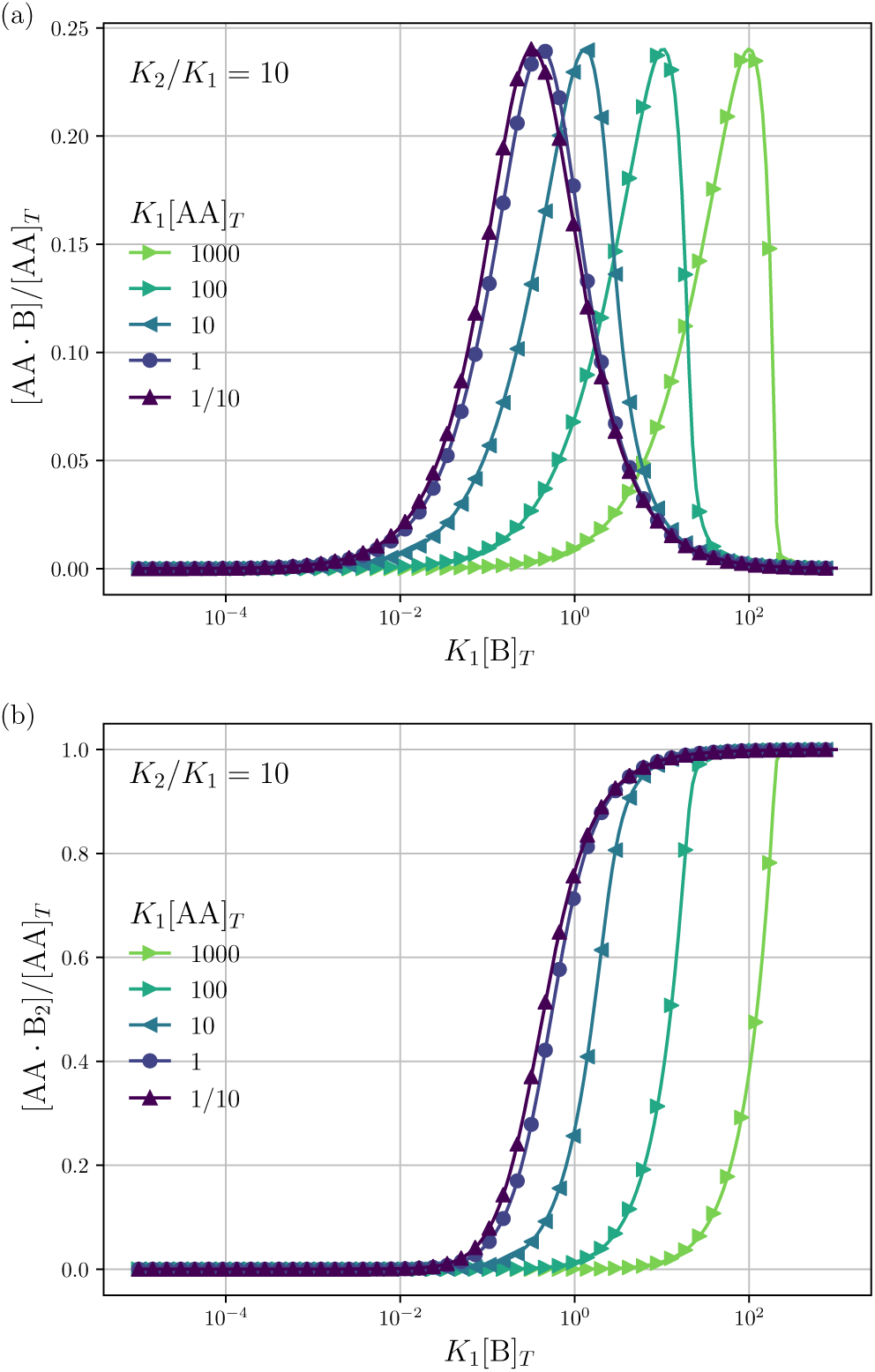
Theoretical predictions for a titration experiment. We show [AA · B]/[AA]_*T*_ (a), [AA · B_2_]/[AA]_*T*_ (b) as a function of *K*_1_[B]_*T*_ for *α* ≡ *K*_2_/*K*_1_ = 10 and several *K*_1_[AA]_*T*_.

## IV. DISCUSSION

Three points of discussion concerning our main Eqs. (6) and (7) are warranted. First, instead of deriving the cubic Eq. (7) for *x*_3_ from Eqs. (2) and (3), we may just as well have isolated *x*_2_ = [B_2_]/[AA]_*T*_. Indeed, cubic expressions for *x*_2_ were reported in Eq. (S) of Ref. [12] and Eq. (25) of Ref. [26] [which we rederive in Appendix F]. However, neither of those articles discussed the dependence of [AA · B] and [AA · B_2_] on the parameter *K*_1_, *K*_2_, [AA]_*T*_, and [B]_*T*_ in much detail.

Second, to model antibody binding to surface-bound antigens, Refs. [11, 12] expressed concentrations of antigens and (partly) bound complexes in numbers per unit area [12, 26]. We note, however, that the governing equations of Refs. [12, 26] could also be cast into the form of Eqs. (2) and (3), that is, with volumetric concentrations *only*, and the effect of reduced positional freedom of surface-bound molecules absorbed into the constants *K*_1_ and *K*_2_. Conversely, though volumetric concentrations appear in our Eqs. (2) and (3), this set of equations can just as well describe a binding process wherein either AA or B is confined to a thin (membrane) surface (see also page 13 of Ref. [1]). Next, Refs. [11, 12] postulated specific relations between *K*_2_ and *K*_1_. Here, we studied Eqs. (2) and (3) for general *K*_1_ and *K*_2_ instead. Hence, in order to apply our mathematical framework to specific binding reactions, one should first determine *α* = *K*_2_/*K*_1_, for example, with the methods of Refs. [17, 18].

Third, the constraint of particle conservation in homo-bivalent ligand-monovalent receptor binding—described in this article—can be especially relevant in cellular contexts, where few molecules of either species may be present. However, for tiny systems with small numbers of particles, the reaction rate equation-type modelling that underlies our results breaks down. One should then account for stochasticity [33], possibly using our continuum results as a benchmark.

## V. CONCLUSION

We have laid out a unified description of the reversible binding of a bivalent ligand to two identical monovalent proteins. The same process has been studied previously, but only in concentration limits of either much more ligands than proteins or vice versa. We have described the binding process for any concentration of ligands and proteins. Comparable concentrations of species can occur both in *in vivo* and in synthetic biological systems. Our theoretical work is built on classical reaction-rate equations. At steady state these reduce to four coupled equations for the concentrations [AA], [B], [AA · B], and [AA · B_2_] of unbound, partly bound, and fully bound protein-ligand complexes, with dependence on four parameters *K*_1_, *K*_2_, [AA]_*T*_, and [B]_*T*_. Only in the limits *ξ* = [B]_*T*_ /[AA]_*T*_ → ∞ and *ξ* → 0 do we recover the results of [Hunter and Anderson, Angewandte Chemie International Edition **48**, 7488 (2009)] and of [Perelson and DeLisi, Mathematical Biosciences **48**, 71 (1980)]; at finite *ξ*, their results contain errors of *O*(*ξ*^−1^) and *O*(*ξ*^3^), respectively. As we move away from these two limits, the concentrations [AA · B] and [AA · B_2_] exhibit a rich and nontrivial dependence on *K*_1_[B]_*T*_. For example, we showed how [AA · B_2_] transition from the bell-shaped crosslinking curves of Ref. [29] to sigmoidal shapes, and how intermediates [AA · B] persist even at high *K*_1_[B]_*T*_.

Our work can be a stepping stone to study the effect of nontrivial protein-to-ligand ratios on hetero bivalent interactions [14, 34–36]. Future work could include how comparable molecular concentrations affect the competition between monovalent and divalent receptors for divalent ligands [28, 37].

## ACKNOWLEDGMENTS

We thank Susanne Liese and Kay Schink for stimulating discussions. MJ and HS were supported by an Advanced Grant from the European Research Council (no. 788954). The research leading to these results has received funding from the European Union’s Horizon 2020 research and innovation programme under the Marie Sklodowska-Curie grant agreement No 801133.

## Appendix A: Derivation of Eq. (2) from reaction rate equations

We repeat Eq. (1)

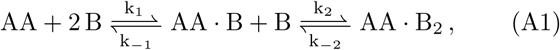

where now *k*_1_ and *k*_2_ and *k*_−1_ and *k*_−2_ are forward and backward reaction rates, respectively. From the law of mass action follow the reaction-rate equations,

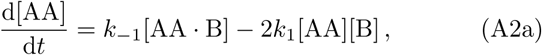

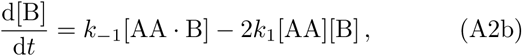

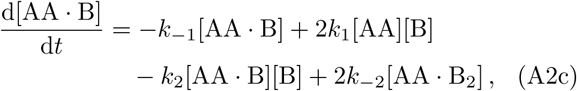

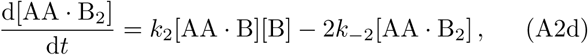

which need to be supplement with initial concentrations of the four species, which we choose as

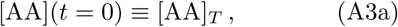

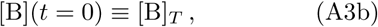

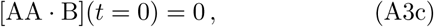

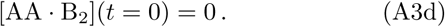

Time-dependent concentrations were studied in [13]. Here, we focus on the steady state, for which Eqs. (A2a) and (A2b) are identical and Eq. (A2c) is the sum of Eqs. (A2a) and (A2d). Writing *K*_1_ = *k*_1_/*k*_−1_ and *K*_2_ = *k*_2_/*k*_−2_ and considering the steady state, we arrive at Eq. (2) of the main text.

## Appendix B: General solution to Eq. (7)

Substituting *x*_3_ = *u a*/3 into Eq. (7) yields the depressed cubic

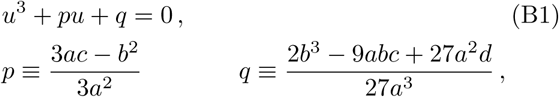

whose solution, with Viéte’s formula, reads

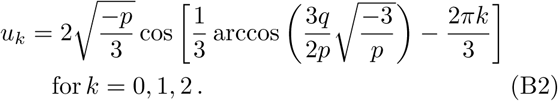

Depending on the values of *κ*_1_, *κ*_2_ and *ξ*, the determinant Δ = −(4*p*^3^ + 27*q*^2^) can be both positive and negative. Hence, for different parameter settings, Eq. (7) has either three real roots or one real and two complex roots.

## Appendix C: Few divalent ligands [AA]_*T*_ ≪ [B]_*T*_ (*ξ* ≫ 1)

For *ξ* ≫ 1, Eq. (7) reduces to

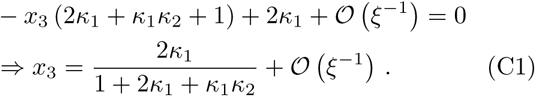

Inserting Eq. (C1) into Eq. (6) and again taking *ξ* ≫ 1, we find

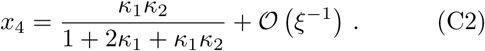

The total receptor occupancy is found as

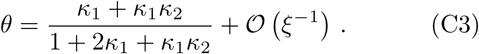

Equations (C1) and (C2) coincide with Eqs. (S20) and (S21) of Ref. [19], which are used draw Fig. 4 therein. From Eq. (C3) we can find the Hill coefficient *n*_*H*_ in terms of the cooperativity parameter *α*,

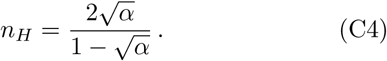

## Appendix D: Few monovalent receptors [AA]_*T*_ ≫ [B]_*T*_ (*ξ* < ≪ 1)

Next, we seek approximate solutions to Eq. (7I:) for *ξ* ≪ 1. Accordingly, we insert the power series 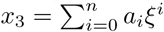 into Eq. (7), collect terms of equal order in *ξ*, and demand the coefficient of each successive order in *ξ* to be zero. For *n* = 3, we find

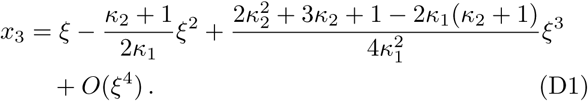

We insert Eq. (D1) into Eq. (6) and find

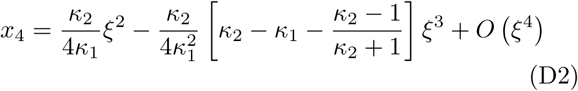

Reference [29] attacked the same problem differently. They stated that [AA] ≈ [AA]_*T*_ can be assumed if [AA]_*T*_ ≫ [B]_*T*_. Then, the term (1 − *x*_3_ − *x*_4_) in Eq. (5a), which stems from [AA] should be replaced by 1, yielding

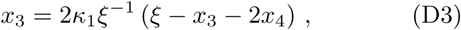

instead. Inserting *x*_4_ [Eq. (6)] as before now yields

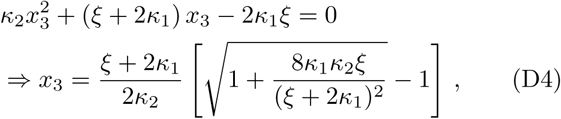

equivalent to Eq. (19) of Ref. [29].

We insert Eq. (D4) into Eq. (6) and find

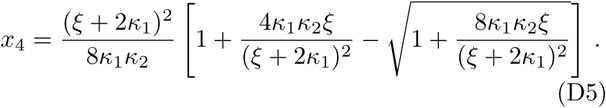

equivalent to Eq. (20) of Ref. [29].

As Eqs. (D4) and (D5) were derived setting *x*_1_= 1, argued on the basis of *ξ* ≪ 1, the *ξ*-range of validity of these expression is not obvious. Expanding Eq. (D4) for small *ξ*,

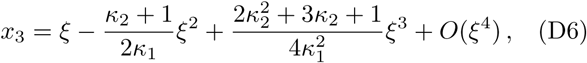

we see that Eqs. (D1) and (D6) differ at *O*(*ξ*^3^). Practically, setting *κ*_1_ = *κ*_2_ = 10^2^, the two approximations Eqs. (D1) and (D6) differ from the numerically found root by 0.0001% and 0.52% at *ξ* = 0.1 and 0.5% and 50% at *ξ* = 1, respectively. As expected: at smaller *ξ*, both approximations are very decent. For *ξ* ∼ 1, Eq. (D1) performs better.

Likewise, expanding Eq. (D5) for small *ξ* yields

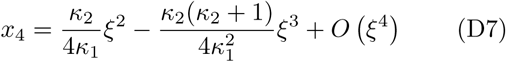

Again, differences between Eqs. (D2) and (D7) appear at *O*(*ξ*^3^). Concluding, Eqs. (19) and (20) of Ref. [29], contain errors of *O*(*ξ*^3^).

## Appendix E: No cooperativity, *α* = *κ*_2_/*κ*_1_ = 1

In absence of cooperativity (*K*_1_ = *K*_2_) we have that *κ*_1_ = *κ*_2_ ≡ *κ* and Eq. (7) simplifies to

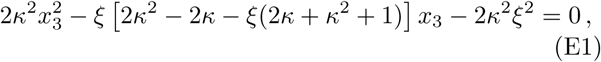

The quadratic Eq. (E1) is solved by

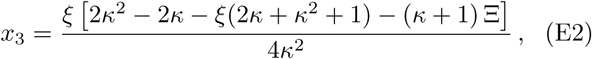

With

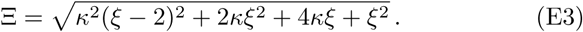

For the interpretation of the plateaus at *κ* ≫ 1 in Fig. 3(a), we note that, for *κ* ≫ 1 and *ξ* ∼ 1,

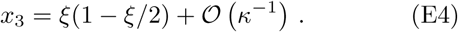

Equation (E4) breaks down for *ξ* > 2, as *x*_3_ > 0 is required.

Inserting Eq. (E2) into *x*_4_ [Eq. (6)] and *θ* ≡ *x*_3_/3 + *x*_4_ yields

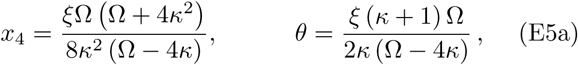

where Ω is defined as

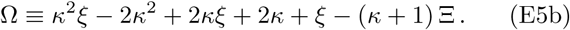

From Eq. (E5a) we find (with sympy)

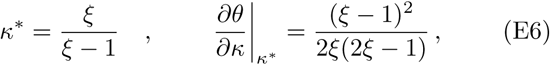

hence

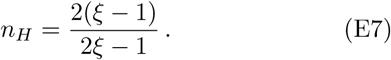

This expression only holds for *ξ* > 1. For *ξ* < 1, *θ* < 1/2; hence, the condition of half occupancy in the definition of the Hill coefficient is never fulfilled.

## Appendix F: Cubic equation for *x*_2_

In terms of the dimensionless parameters (and *x*_1_ = [AA]/[AA]_*T*_ and *x*_2_ = [B]/[AA]_*T*_), Eqs. (2) and (3) read

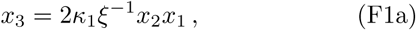

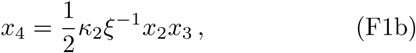

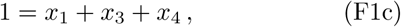

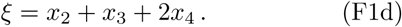

From Eqs. (F1c) and (F1d) we find

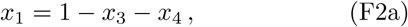

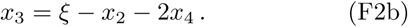

From Eqs. (F1b) and (F2b) we find

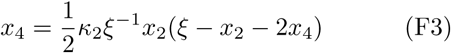

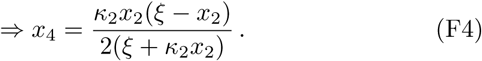

Inserting Eqs. (F2a) and (F2b) into Eq. (F1a) we find

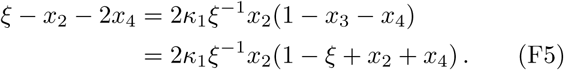

Inserting Eq. (F4) gives

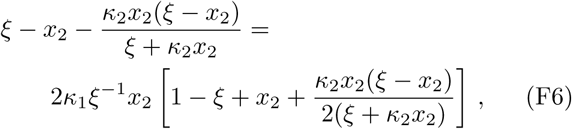

which yields

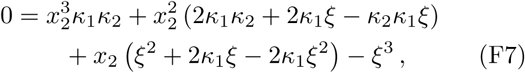

or, in our original notation,

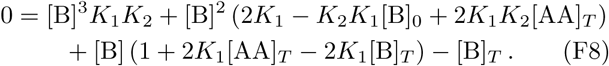

With Eqs. (F7) and (F8), the original problem formulation of four coupled equations in Eqs. (2) and (3) has been reduced to a single cubic equation for [B]. As such, it forms an alternative to the cubic equation for [AA · B] in Eq. (7) of the main text. Equation (F8) is equivalent to Eq. (25) of Ref. [26]—up to factor 2 discrepancies in a few places, which we trace back to her Eq. (15), the counterpart of our Eqs. (F1a) and (F1b), which does not include prefactors 2 and 1/2. Redefining our *K*_1_ → *K*_1_/2 and *K*_2_ → 2*K*_2_ lifts these discrepancies. Moreover, Eq. (F8) is equivalent to Eq. (S) of Ref. [12] in the case that their “nonreactive fraction parameter” *nr* is set to *nr* = 0.

